# Biophysical constraints on avian adaptation and diversification

**DOI:** 10.1101/2023.10.26.564103

**Authors:** Ferran Sayol, Bouwe Reijenga, Joseph A. Tobias, Alex L. Pigot

## Abstract

The capacity of organisms to adapt to vacant niches or changing environments is limited by physical constraints on morphological evolution. Substantial progress has been made in identifying how these constraints shape the form and function of producers (plants), but our understanding of evolutionary limits in consumers (animals) remains highly limited, in part because the requisite data have not been available at sufficient scale. Using morphometric measurements for all birds, we demonstrate that observed variation is highly restricted—both for beak shape and overall body shape—to triangular regions of morphospace with clearly defined boundaries and vertices. By combining morphometric data with new information on physical functions of measured traits, we provide evidence that the extent of avian morphospace is constrained by biophysical trade-offs between three functional objectives (strength, reach and engulfment capacity) that characterize resource acquisition and processing by the beak, and three locomotory modalities (aerial, aquatic and terrestrial) that characterize avian lifestyles. Our results suggest that over avian evolutionary history, trajectories of morphological change trend towards the vertices, with birds evolving from a core of biophysical generalists to biophysical specialists, associated with faster macroevolutionary turnover of lineages at the periphery or morphospace. Our analyses reveal that the structure of avian morphological diversity follows relatively simple rules defined by biophysical constraints and trade-offs, shedding light on the process shaping modern animal diversity and responses to environmental change.

## Main text

Over billions of years of evolution, life has explored an enormous diversity of physical forms. This morphological diversity largely reflects the variety of different strategies employed by organisms for capturing and allocating resources and for exploiting environments with radically different physical states and properties (1–5). Yet, this morphological diversity often seems highly constrained in comparison to all geometrically possible options (6, 7). Ultimately, all organisms must abide by the universal laws of physics (8, 9), imposing bounds on the range and combination of traits that are biophysically possible given a particular body plan and that are viable in a world of relentless competition and selection (10, 11). Understanding the strength, nature and identity of these constraints is essential not only for explaining the evolution of phenotypic diversity within and across species (12, 13), but also for predicting the adaptive capacity and resilience of populations under rapid environmental change (14–16). However, despite this fundamental importance, we lack a comprehensive understanding of how the physical environment organises and limits the diversity of morphological forms observed in nature, especially among heterotrophic animals that comprise much of global biodiversity.

Constraints to evolution imposed by the physical environment are expected to lead to regular patterns in the morphological forms, shapes and structures of organisms, that is ‘morphospace’ (17, 18). In particular, because organisms are subject to multiple physical constraints, this results in trade-offs, as the performance of one functional objective comes at the expense of another (7, 19). In this case, natural selection is expected to optimise organism performance across these multiple objectives, eliminating combinations of traits where performance is reduced, and resulting in multiple traits aligning along a limited number of independent morphological dimensions (17). The co-variation between multiple aspects of organism function and body size represents one such axis along which morphological diversity is organised (20, 21). The advent of large-scale datasets characterising whole-organism plant form have identified additional axes of constraint acting on photosynthesising primary producers, providing an organising framework for understanding and predicting how autotrophic species and communities respond to environmental change (1, 22, 23). In contrast, due to a lack of quantitative morphological data, the search for the key constraints and trade-offs shaping the evolution of morphological diversity across higher trophic levels has lagged far behind, despite the critical role of these organisms in driving ecosystems processes and supporting plant diversity (24, 25).

Here we test for constraints on the morphospace occupied by birds, which despite conservatism in their body plan have radiated across multiple trophic levels and physical environments (2, 26). Exceptionally, owing to recent efforts in digitizing museum specimens, quantitative data is now comprehensively available across a number of key morphological traits allowing us to characterise the size and shape of morphospace occupied by almost all ∼10,000 extant bird species (27). We initially focus on variation in the beak, the primary anatomical apparatus used by birds for resource acquisition and processing and a model system for understanding the functional, developmental and genetic basis of trait variation (28–30). Using linear measurements of beak length, width and depth, we first characterise the volume and geometry of beak morphospace. We demonstrate that beaks occupy a highly restricted volume of what is geometrically possible, with variation in beak shape largely restricted to a triangular region of morphospace that reflects trade-offs among three primary functional objectives required for acquiring and processing resources. Using measurements of the tail, wing and tarsus, we show that variation in bird body shape is similarly restricted to a triangular region of morphospace, reflecting constraints imposed by the three physical states of matter through or upon which birds move. We show how these biophysical constraints on beak and body form provide a potential framework for understanding the evolution of bird morphological diversity and species resilience to global environmental change.

### Global beak morphospace

Based on log-transformed linear measurements of beak length, width and depth we used the convex hull enclosing all species to calculate the volume and shape of 3-d beak morphospace occupied by birds (**Figure 1a**). We compared this to a series of different null models that make contrasting assumptions about how species could be distributed throughout beak morphospace. Null model 1 assumes that trade-offs are absent and that within the observed extremes any combination of traits is equally viable. This leads to a morphospace resembling a cube (**Figure 1b**). We found that real beaks occupy a volume of morphospace much smaller than expected under this null model (14% of volume occupied by null model 1) (**Supplementary Figure 1a**), indicating that coordination among beak traits and/or selection against extreme trait combinations (i.e. the minimum or maximum values of multiple traits in the same species) limits the variety of beak forms. Coordination among traits is clearly the major factor because, despite exhibiting more extreme trait values, the volume of empirical morphospace is also substantially smaller (27% of volume occupied by null model 2) than expected under an alternative null model (Null model 2) in which traits vary independently but where extreme trait combinations are not permitted (**Figure 1c, Supplementary Figure 1a**). In other words, trade-offs among traits, rather than restrictions on extreme trait combinations *per se*, appears to be the primary way in which observed variation in beak form is constrained below what is geometrically possible.

**Fig. 1.**
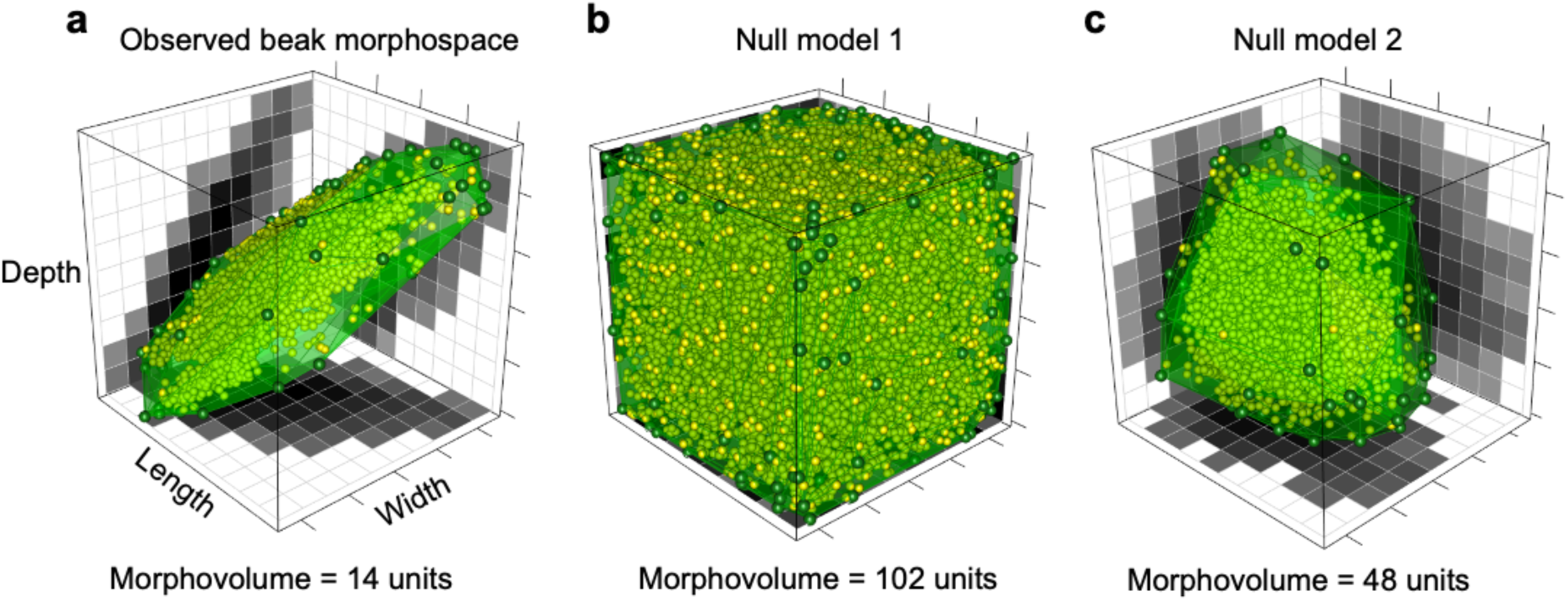
The volume and shape of avian beak morphospace. Points show species (*n* = 9,912) in 3D beak space defined by beak length, width and depth measurements. Green polygons show the boundary of the convex hull enclosing all species, with dark green points indicating the vertices of the hull. The density of species is projected onto each 2-dimensional plane. Dark shading indicates a higher density. (**a**) empirical beak morphospace. (**b-c**), alternative null models of morphospace occupation. (**b**) Null Model 1 assumes there are no trade-offs among beak trait dimensions or selection against extreme beak forms. (**c**) Null Model 2 assumes beak trait dimensions vary independently but extreme beak forms are not viable. The volume of morphospace occupied by real beaks and each null model are indicated.

A principal component (PC) analysis indicates that the main dimension of beak form variation represents differences in size, capturing 86% of the variance in beak dimensions (**Supplementary Table 1**). Beak size varies by >5 orders of magnitude across species (from the Andean palm swift (*Tachornis phoenicobia*) to the Shoebill (*Balaeniceps rex*)), likely reflecting the enormous variation in the size of exploited food items (2, 30), as well as additional selective pressures (e.g. thermoregulation (31, 32)). Each trait has an almost equivalent loading on PC1 indicating that beak shape is largely invariant with size (**Supplementary Table 1**), so that the most massive beaks are essentially scaled up versions of the smallest beaks (**Figure 1a**). As a result, given the observed bounds of each beak trait, bird beaks have evolved to occupy almost the full range of sizes (84%), from the most massive to minute, of what is geometrically possible (**Figure 1b, Supplementary Figure 1b**). Taken together, these results reveal how the restricted volume of beak morphospace largely reflects coordination among beak traits that results in a lack of extreme shapes rather than simply restrictions on size (**Supplementary Figure 1**). To examine constraints on the beak independent of size, in our subsequent analysis we therefore used variation in the second and third dimensions of beak PC space, which describe the relative length (PC2) and relative width and depth (PC3) of the beak respectively.

### Geometry of beak shape variation

Rather than occupying an amorphous region across the 2-d plane defined by PC2 and PC3, beak shape morphospace resembles a triangle, with diffuse but relatively clearly defined edges and vertices (**Figure 2a**). Morphospaces conforming to simple geometrical patterns, like lines, triangles or trapezoids, are predicted by optimisation theory when there are trade-offs to performing a limited number of functional objectives (17, 33). According to this theory, performance of a given functional objective is optimised at a single point in morphospace, and declines away from this ‘archetype’. When selection acts on the performance of two objectives, trade-offs will cause species traits to be organised along the line connecting the two archetypes. This region where performance is maximised is termed the Pareto front. With trade-offs among three functional objectives, the Pareto front expands to occupy a triangular region of morphospace, with each vertex of the triangle corresponding to realised or theoretical trait combinations that are specialized to perform one of the three functional objectives (17). With a large number of functional objectives, morphospace would increasingly resemble a circle. Thus, according to this theory, the number of vertices defining morphospace is indicative of the number of functional trade-offs shaping morphological diversity.

**Fig. 2.**
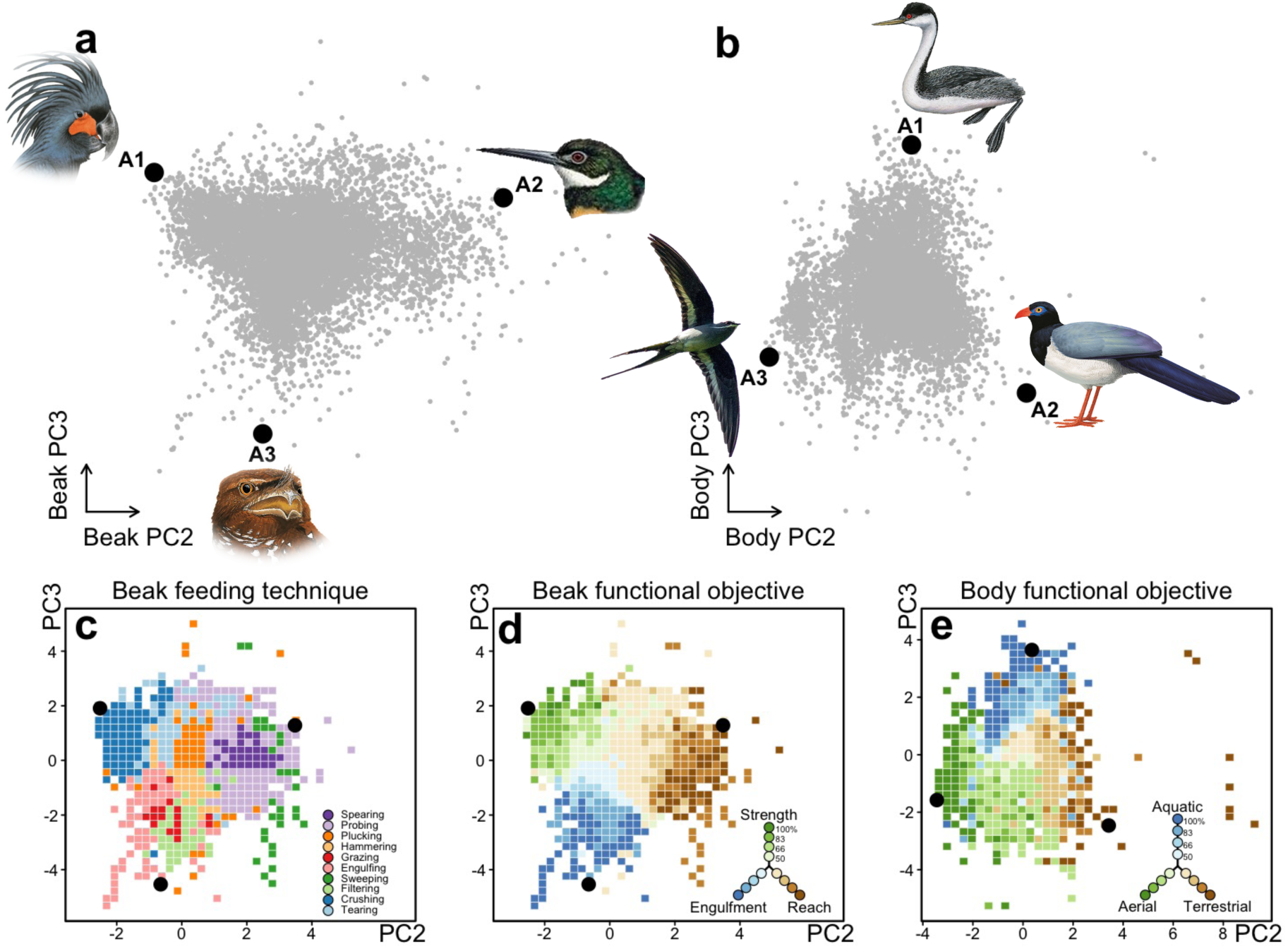
Trade-offs constrain beak and body shape variation to simple triangles (*n* = 9,912 species). (**a**) Beak PC2 and PC3 describe variation in relative beak length (PC2) and relative width and depth (PC3). (**b**) Body PC2 and PC3 describe variation in relative length of the legs versus wings (PC2) and the tail (PC3). Green points show individual species (*n* = 9,912). Darker shading indicates a higher local density of species in morphospace. Black circles show the position of ‘archetypes’ (i.e. vertices) defining the best fitting triangle for describing beak and body shape variation. Examples of species close to archetypal positions are illustrated. (**c**) shows the dominant feeding technique (*n* = 10 techniques) mapped across beak morphospace and (**d-e**) the dominant functional objective mapped across (**d**) beak (strength, reach and engulfment capacity) and (**e**) body morphospace (aquatic, terrestrial, aerial).

To formally test how well beak shape morphospace conforms to a triangle, we fit to the boundaries of the empirical point cloud a series of polygons, varying both the number and location of vertices. When polygons are allowed to take irregular shapes, adding more vertices (i.e. archetypes) will always improve fit. However, we found that the improvement in fit beyond three vertices is marginal (**Supplementary Figure 2**). For regular polygons, adding more than three vertices results in a poorer fit (**Supplementary Figure 3**), confirming the visual impression that beak shape morphospace can be well described by a triangle. The first archetype (A1) of beak shape morphospace describes beaks that are short, narrow and deep (**Figure 2a**). The second archetype (A2) describes beaks that are of intermediate length, flat and wide. The third archetype (A3) describes beaks that are long relative to their width and depth. There is some indication for a possible fourth archetype, corresponding to beaks that are long and deep (**Supplementary Figure 2**). However, the vertex angle is wide and may instead simply be approximating the curved upper edge of morphospace. Such curved edges connecting archetypes are not unexpected under the Pareto front concept, and can arise if the performance of an objective is optimised across a region rather than at point in morphospace or if changes in performance of a given objective do not occur uniformly across morphospace (18). While the restricted volume and geometry of beak morphospace is consistent with the idea that trade-offs among a small number of functional objectives have constrained the variety of beak forms, a stronger test requires identifying these objectives and how specialization in their performance is distributed across beak morphospace.

### Trade-offs constraining beak form and function

Bird beaks provide multiple functions (e.g. nest building (34), thermoregulation (31, 32)) but their primary task is the acquisition and processing of food (2, 35). Species data on the relative use of 10 major feeding techniques (e.g. probing, crushing, engulfing) show how these map onto distinct regions of beak morphospace confirming this tight functional association (**Figure 2c**). However, the number of distinct feeding techniques far exceeds the three functional objectives implied by the triangular shape of beak morphospace. To make sense of this, we hypothesise that the different techniques for acquiring and processing resources can ultimately be collapsed down to the performance of three primary functional objectives, and that it is trade-offs in the performance of these objectives that provide the principal constraints on the variety of realised beak shapes.

Processing of food items requires mechanical strength to prevent breakage of the beak when a force is applied across it (e.g. when tearing or crushing prey) (35). Experimental data on bite force across species show that greater mechanical strength is provided by a larger beak size and a relative deepening of the beak (i.e. low PC2 and high PC3) (36), corresponding to Archetype 1 in the top left region of beak shape morphospace (**Supplementary Figure 4a**). However, a deeper beak comes at the expense of a slower closing velocity (**Supplementary Figure 4b**) and a reduction in the efficiency of prey acquisition (37), which should be maximised by either a relative elongation (i.e., higher PC2) or widening (i.e., lower PC3) of the beak. Elongated beaks provide a longer closing surface and sweeping circle for capturing fast moving prey (35, 38), as well as access to concealed food items, such as animals buried in softs sediments, or seeds and nectar within flowers. We refer to this second objective as ‘reach’ capacity. Alternatively, widening of the beak increases ‘engulfment’ capacity, including the volume of water that can be filtered for plankton (39) or plant material that can be grazed (40), or the probability of capturing small insect prey while in flight (41). Thus, for a given beak size, increases in any of strength, reach or engulfment capacity is expected to come at the expense of performing the other objectives.

Based on information describing the relative use of different feeding techniques and food types (e.g. seeds versus invertebrates) we calculated scores of the relative importance of strength, reach and engulfment capacity for each species (**Supplementary Table 2**). We found that these three functional objectives map out with remarkable fidelity to the three vertices of morphospace, in a way that matches theoretical expectations (**Figure 2d, Supplementary Figure 5**). At each vertex, species are specialized at performing the corresponding functional objective, with the specialization on that objective then declining away from that vertex (**Figure 2d, Supplementary Figure 6**). Species along the boundaries or centre of the morphospace perform two and three of the functional objectives respectively depending on their position in morphospace. This close and one-to-one mapping of objective specialization to position within morphospace supports the idea that trade-offs in the performance of these different functional objectives constrain the set of beak forms that are viable over evolutionary time.

One potential criticism of our analysis is that the simple linear measurements of beak length, width and depth, that we used have failed to capture the primary axes of beak shape variation. We tested this by repeating our analysis on a subset of species (*n* = 2026 species) using shape axes derived from 3-D beak scans that fully capture the complexity of shape, including curvature (30). The two primary axes of shape variation derived from 3-D scans (PC1_scan_ & PC2_scan_) account for the majority (87%) of total beak shape variation and are aligned with our original PC axes based on linear measurements (**Supplementary Table 1)**. Accordingly, all our key conclusions remain unaltered. The filling of beak morphospace across the plane defined by PC1_scan_ & PC2_scan_ also conforms to a triangle (**Supplementary Figure 7**), with vertices corresponding to relatively long, wide or deep beaks and with each of these vertices exhibiting the same match to strength, reach and engulfment capacity (**Supplementary Figure 8**). Thus, the simple geometric structuring of avian beak shape occurs regardless of the way these traits are measured.

### Biophysical constraints on body morphospace

The beak is the primary apparatus used by birds for acquiring and processing resources. However, it is an incomplete description of the trophic niche. Species with similar shaped beaks can utilise substantially different foraging strategies, that describe how species move through the environment to locate and attack their prey. To address this, we extended our analysis to consider constraints on variation in body shape using a PC analysis of the key traits linked to locomotion in birds: the wing, tail and tarsus (**Supplementary Table 3**). We found that the same pattern of constraint identified for beak shape also apply to body shape. Body shape morphospace defined by PC2 and PC3 resembles a triangle and can be well described by a polygon with three vertices (**Figure 2b, Supplementary Figures 2 & 3**). We hypothesised that the primary constraints on bird body shape are imposed by the physical environments in which species forage. Scoring the proportional use of different physical environments confirms this pattern (**Figure 2e, Supplementary Table 4**). We show that the three vertices of the Pareto front correspond to body forms specialized for moving either through water (A1), across terrestrial surfaces (i.e. ground and arboreal) (A2) or through the air (A3), with intermediate forms utilising multiple physical media (**Figure 2e, Supplementary Figure 5**). Thus, similarly to beak shape, our results suggest that the primary axes of variation in body shape are also constrained by trade-offs among a small number of functional objectives, in this case related to three locomotory modalities (aerial, aquatic and terrestrial).

### Evolutionary dynamics of a bounded morphospace

The bounded nature of bird beak and body morphospace inferred by our analysis help explain several macroevolutionary trends. The evolution of beak shape among extant birds is characterised by an early expansion of disparity, while over the last ∼50 million years the volume of beak morphospace has remained relatively stable, despite rapid and ongoing species evolution (30). During this latter period of morphospace infilling, evolutionary convergence has been pervasive, with distantly related bird lineages repeatedly converging on the same regions of beak and body morphospace (2, 42). The constraints on morphospace that we identify could explain both the stalling of beak disparification as well as the widespread evolutionary convergence in morphology as divergence to occupy novel regions of morphospace is restricted.

To explore how bird morphospace has filled over time, we performed a phylogenetic reconstruction of beak and body shape, analysing the macroevolutionary dynamics in the direction of phenotypic evolution. Our results show a net flow of lineages from the densely packed core of morphospace to the periphery and corners of the Pareto front (**Figure 3 a,d**). A net outward flow of lineages through morphospace is expected under a simple model of diffusion. However, the observed dynamic is stronger, with twice as many lineages inferred to evolve outwards to the periphery compared to inwards to the centre (**Supplementary Figure 9**). These results support the hypothesis that lineages become more biophysically specialized over macroevolutionary time (43, 44). However, this pattern should be interpreted with caution given the known challenges of inferring long extinct ancestral forms (45). This net outward flow of lineages also does not explain constraints on morphospace, and why we observe more species in the core compared to the edges of the Pareto front(2, 46, 47).

**Fig. 3.**
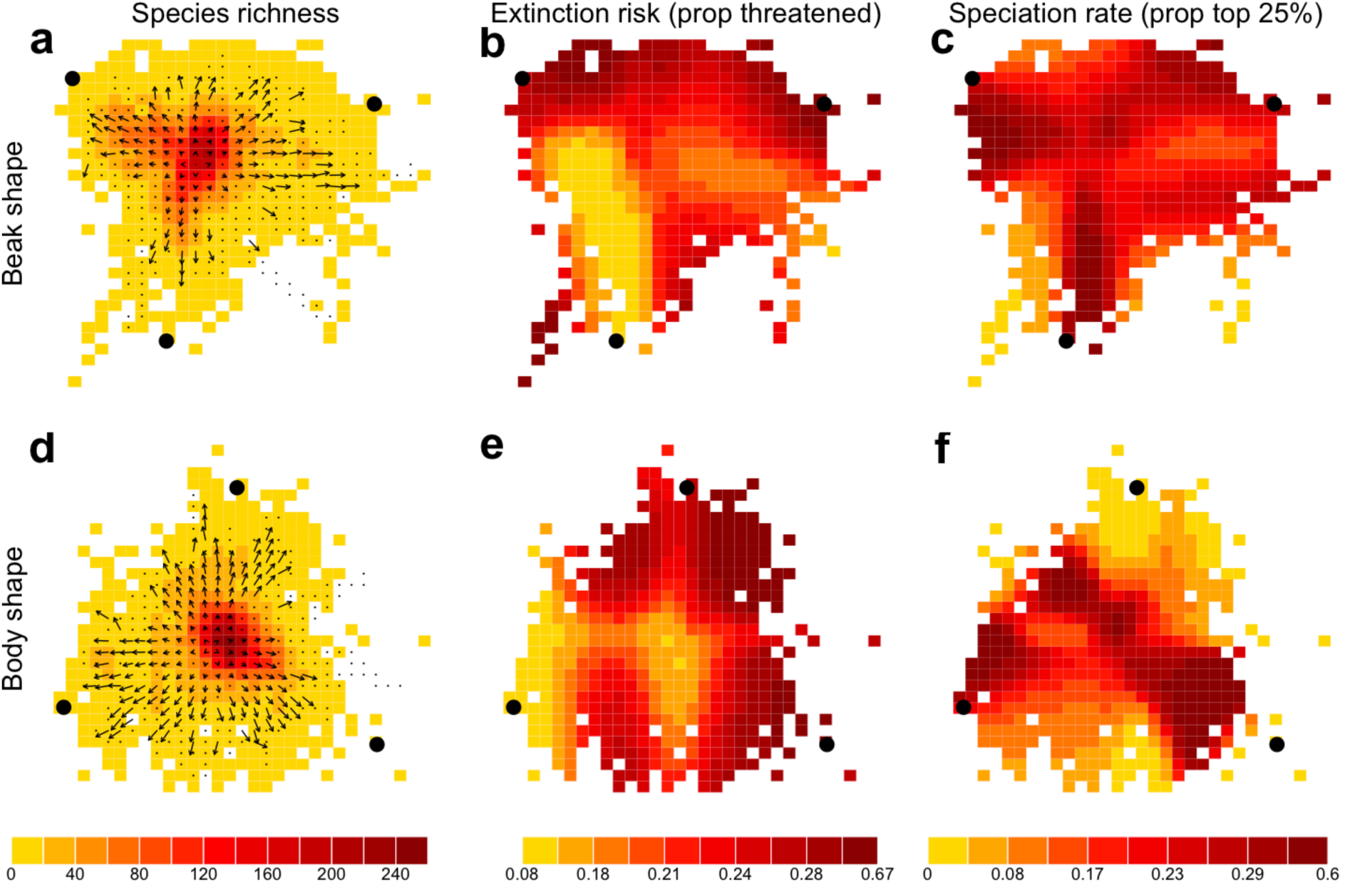
Macroevolutionary dynamics of beak and body shape morphospace. (**a, d**) Species richness (colors) and the mean direction of trait evolution (arrows) mapped across beak and body shape morphospace. Predicted probability of a species being threatened with extinction (**b, e**) and being in the top speciation rate quartile (**c, f**) mapped across beak and body morphospace. Archetypes defining the triangular Pareto front are indicated as black points. In (**a, b**) arrows show the mean direction of evolution along the branches reconstructed to intersect each grid cell. Longer vectors indicate a greater consistency in direction (i.e. a lower variance). Arrows are replaced with points if the consistency in the direction of evolution was no greater than expected under a model of random trait diffusion.

One possible explanation for the relative scarcity of species near and beyond the edges of the Pareto front might be a higher risk of extinction of biophysically specialized species (46, 48–50), lower rates of speciation or both (51). Mapping the incidence of currently threatened species and of lineages with historically fast speciation rates, reveals a number of extinction (**Figure 3 b,e**) and speciation (**Figure 3 c,f**) hotspots across beak and body morphospace. These hotspots of high extinction risk and speciation rate tend to cluster around the edges, and often near the corners of the Pareto front (i.e. archetypes) where biophysical specialization is highest, albeit rarely at the precise location of the inferred archetypes. The occurrence of these extinction and speciation hotspots at the edge of the Pareto front could indicate a dynamic of greater macroevolutionary turnover of lineages, which by opening new ecological opportunities could drive the net outward flow of lineages from the core, while also capping the density of species at the edge. However, this suggestion remains speculative. Some peripheral regions of morphospace instead coincide with extinction (e.g. Body A3) or speciation (e.g. Body A1) coldspots (**Figure 3 c, f**). Furthermore, in species level models accounting for phylogenetic non-independence and other potential confounding variables, we found that there was no consistent effect of distance to archetypes on speciation rates, with extinction risk only significantly increasing towards body, and not beak, archetypes (**Supplementary Figure 10 and Supplementary Table 4**). Thus, while the distribution of current threat across morphospace is in line with the notion that specialized species are more sensitive to anthropogenic environmental change(52, 53), our global analysis suggests that biophysical specialization alone does not consistently predict the propensity of lineages to split or persist.

A potential explanation for the inconsistency of these findings is that a focus on current patterns of threat and recent rates of speciation may fail to capture the macroevolutionary dynamics that have shaped morphospace throughout avian history (54, 55). Alternatively, if the lack of a general link between diversification dynamics and biophysical specialization does also operate over deep macroevolutionary time, this would suggest that the triangular boundary of avian beak and body morphospace is not maintained by selection at or above the species level (56), but instead reflects microevolutionary processes, with constraints on adaptation inhibiting the evolution of more extreme trait combinations. The edges to morphospace that these constraints impose, need not correspond to ‘hard’ boundaries, beyond which trait combinations are biophysically impossible (8, 9). Afterall, the peripheral regions of trait space beyond the Pareto front although sparsely filled are not completely devoid of species. Instead, we suggest that while trade-offs constrain most species to occur within the triangular Pareto front(17, 18), a relaxation of, or escape from, these trade-offs due perhaps to geographical isolation from competitors or key innovations(57, 58), allows some lineages to persist beyond the Pareto front. In any case, release from these constraints appears to be sufficiently rare or fleeting so as not to disrupt the overall triangular structure of morphospace.

While trade-offs in the performance of a limited number of biophysical objectives appear to constrain the shape of avian morphospace, ecological constraints on adaptation could drive variation in species density within these bounds. The central core of high species density (**Figure 3a, b**), is largely comprised of arboreal gleaning invertivores for which the high structural complexity of their predominantly forest habitat and the enormous diversity of invertebrate prey could lead to a finer partitioning of ecological niches thus enhancing species coexistence(35),(59, 60). Our data support this explanation, showing that species near the centre of morphospace have narrower foraging niches consistent with the compression of niche breadth linked to high species packing (**Supplementary Figure 11**). Our results further show that this ecological niche specialization does not correspond to biophysical specialization, and that arboreal gleaning invertivores–which comprise by the most speciose foraging niche–are actually biophysical generalists (**Supplementary Figure 11**), with beak shapes selected for a trade-off between reach, engulfment and strength capacity, and with body shapes reflecting a trade-off between terrestrial and aerial locomotory modalities.

The morphology of birds—with their wings and keratinous beaks—is unique in the animal kingdom, but occupies only a small subset of the enormous morphological variety of heterotrophic organisms. The limits to avian evolution inferred by our analysis would thus reflect the biophysical constraints operating in the context of the developmental pathways and body plan of birds. Yet, these trade-offs would also seem sufficiently fundamental to suggest a general framework for organising this wider animal diversity. For evolutionary radiations spanning multiple realms, our findings predict that the primary axes of body shape variation should conform to lines, triangles or quadrilaterals according to the number of physical states of matter through or upon which organisms move (61, 62). Similarly, trade-offs between strength, reach and engulfment capacity that constrain beak evolution to a triangular Pareto front could apply to the skulls of other vertebrates that also primarily catch and process resources using their mouths (38, 63). An insectivorous nightjar (**Figure 2a, A3**) swooping through a forest, its mouth agape, draws parallels with a lunging baleen whale (39), while the deep beak of a parrot (**Figure 2a, A1**) breaking the hard kernel of a seed, resembles in its function the robust skull of a bone cracking hyena (64). A butterfly plucked in mid-flight by the tweezer like beak of a jacamar (**Figure 2a, A2**), meets the same fate as a fish snagged by a gharial as it sweeps its elongated snout through the water (38). While the diversity of body plans and lack of quantitative morphological data has previously hindered efforts to integrate and combine morphospaces spanning multiple animal classes, the identification of universal biophysical constraints can allow us orientate and position species across these trade-offs, moving us closer to a general and mechanistic understanding of the origins, maintenance and functioning of animal biodiversity and how this responds to environmental change.

## Methods

### Morphological data

We obtained linear measurements for morphological traits related to feeding and locomotion for *n* = 9,993 species (all known birds according to BirdTree taxonomy (65)) from the AVONET dataset(27). We selected data for seven traits: beak length, width and depth, tarsus length, tail length, wing chord and Kipp’s distance (measured as the distance between the wingtip and the tip of the first secondary feather). After excluding species with missing information in some of the target traits, we had a sample of *n* = 9,912 species. Traits were measured using a consistent protocol for 89,963 individuals (mean per species = 9.00) using museum specimens and individuals collected in the field. Most of the variation in traits occurs among (98.7 %) rather than within (1.7 %) species, justifying the use of species mean trait scores (27). For the beak and body separately, we summarized the variation in morphology using a principal components (PC) analysis, following log-transformation and z-scaling of the trait variables. For body shape, species where Kipp’s distance was close to zero (values = 0.1, n = 6 species) were excluded from the analysis. We selected the principal components representing shape (i.e. PC2 and PC3) and rescaled these to unit variance and a mean of zero prior to further analysis. Trait loadings for each beak and body trait are provided in **Supplementary Table 1 and Supplementary Table 3**, respectively. Measures of beak length, width and depth capture relative differences in beak dimensions but effectively assume a conical shaped beak. To ensure our results were not artefacts or treating beak shape in this simplistic way, we repeated our analysis using the first and second PC axis from beak shape measurements for *n* = 2028 species based on a Procrustes superimposition of landmarked 3D beak scans (30). These measurements capture the full complexity of shape, including curvature.

### Beak feeding techniques

To quantify the beak feeding techniques and functional objectives employed by each species, we developed a new dataset describing the % used of 10 broad feeding techniques utilised by birds: plucking, crushing, tearing, hammering, spearing, probing, sweeping, filtering, grazing and engulfing. While not exhaustive, our choice of categories reflects the level of detail typically available in standard textual descriptions of species diets and foraging behaviour and correspond to general strategies utilised across distinct ecological niches. A detailed description of each feeding technique is provided in **Supplementary Table 7**.

To efficiently score the % use of each beak feeding technique for all ∼10,000 birds, we performed a multi-step approach. Because species morphology and ecology are often relatively strongly conserved, we divided our dataset into taxonomic families (*n* = 194 families). For each family, we used textual descriptions in the literature to identify the feeding techniques corresponding to each foraging niche utilised by the species in that family. For this, we used a existing global database(2) that provides species level scores of the relative use of 32 different foraging niche categories, each of which describes a combination of resource type (e.g. ‘invertebrate’) and foraging behaviour (e.g. ‘ground gleaning’) (**Supplementary Table 6**). For example, the ‘ground gleaning invertivore’ foraging niche is utilised by (among many other families) the Kiwi’s (*Apterygidae*) and Pheasants (*Phasianidae*), but the species in these two families utilise different feeding techniques, ‘probing’ and ‘plucking’ respectively, when foraging for invertebrates on the ground. We note that while all species within a given family will receive the same beak feeding technique score for a particular foraging niche, the total use of a given beak feeding technique can still vary across species within that family due to differences among species in the utilisation of each foraging niche. For example, while all hummingbirds (*Trochilidae*) are scored as utilising the ‘probing’ feeding technique when foraging for nectar, the % contribution of ‘probing’ to a species’ diet will vary as some species also obtain resources through other foraging niches (e.g. ‘aerial screening invertivore’) which involve different feeding techniques (e.g. ‘plucking’). For a subset of 14 families, where different species were obviously utilising substantially different beak feeding techniques within a single foraging niche, we scored the genera and/or species within that family individually.

### Beak functional objectives

To quantify the importance of the three beak functional objectives for each species, we first assigned the % importance of each of these objectives for each beak feeding technique (**Supplementary Table 2)**. The quantitative information needed to do this in a precise way is lacking, but based on the extensive literature on beak biomechanics and animal feeding behaviour we can at least coarsely provide an indication of the relative importance of each objective for each beak feeding technique. For example, ‘crushing’ is a technique employed for breaking open hard resources such as shells, seeds and nuts, and was assigned a score of 1 for ‘strength capacity’ (and 0 for ‘reach’ and ‘volume’). In contrast, ‘probing’, which involves the relatively slow or gentle insertion of the beak into a crevice or sediment to extract food, received a score of 1 for ‘reach capacity’. Some beak feeding techniques, however, can be assigned to more than one functional objective. For instance, the ‘hammering’ technique was assigned a score of 0.5 for ‘strength capacity’ and 0.5 for ‘reach capacity’. With these scores, the importance of each functional objective to each species was calculated by multiplying the importance score for a beak feeding technique by the % use of that technique.

### Body functional objectives

Quantifying the importance of different body locomotory objectives (Aerial, Aquatic and Terrestrial movement) for each species is challenging at a global scale. Except for the very small number of flightless species (n = 60 species) (66), almost all birds make use of aerial movement during foraging or migration, but quantitative data on the time a bird spends in flight, or indeed on the ground and in/on the water, is not widely available. Therefore, we developed a simple system for translating our existing species level database^2^ describing the relative use of 32 foraging niche categories into scores that capture the importance of aerial, aquatic or terrestrial environments during feeding. For instance, foraging niches that are restricted to the air (e.g. ‘Aerial screening’) received an ‘Aerial’ score of 1. In contrast, foraging niches that involve searching for food entirely on the ground (e.g. ‘Invertivore glean ground’ or ‘Herbivore ground’) received a ‘Terrestrial’ score of 1, whereas foraging niches that only involve swimming or diving during feeding, received an ‘Aquatic’ score of 1. Some foraging niches require optimising across multiple locomotory objectives. For instance, the arboreal gleaning foraging niche can be considered to use both a terrestrial locomotion when perching and an aerial locomotion when flying among branches, and hence is assigned a score of 0.5 for ‘Aerial’ and 0.5 for ‘Terrestrial’. Similarly, species that plunge into the water to catch fish, which require both flying and swimming for their foraging technique, are assigned a score of 0.5 for ‘Aerial’ and 0.5 for ‘Aquatic’ locomotion. The scoring system for translating between foraging niches and locomotory objectives is provided in **Supplementary Table 4**. With these scores, the importance of each objective to each species was calculate by multiplying the importance score for a foraging niche by the % use of that niche by each species.

### Phylogenetic data

To understand the evolutionary dynamics of biophysical specialization and its influence on species capacity to respond to environmental change, we used the time-calibrated molecular BirdTree phylogeny (65) based on the Hackett backbone topology (67). We downloaded a distribution of 100 phylogenetic trees only including species with genetic data (*n* = 6670 species) for use in phylogenetic reconstructions of evolution. We also downloaded a distribution of 100 complete phylogenies, including species inserted on the basis of taxonomy because they lacked genetic data (*n* = 9993 species), for use in modelling how morphological position influences speciation and extinction. We computed the maximum clade credibility (MCC) of the complete phylogeny using the R package ‘phangorn’ (68).

### Null models of beak morphospace volume and shape

We quantified the volume and shape of the 3-d beak morphospace occupied by species defined by log-transformed length (L), width (W) and depth (D) measurements. Morphospace volume was quantified using a minimum convex hull. Having identified size as the major axis of beak variation (PC1), we quantified the beak size of each species as the ‘beak volume’, assuming beaks are conical in shape (beak volume = pi x 1/3 x L x D x W). Throughout we refer to ‘beak volume’ as ‘beak size’ to avoid confusion with beak morphospace volume. We calculate the variation in beak size across species as the difference between the log_10_ transformed minimum and maximum beak size values. This metric captures the order of magnitude difference between the smallest and largest beaks (**Supplementary Figure 1b**).

Following Diaz et al. (1), we compared occupied morphospace volume and the variation in beak size to that expected under null models that make contrasting assumptions about how species are distributed throughout trait space (**Supplementary Figure 1**). Null model 1 assumes that there are no correlations among beak dimensions or trade-offs so that species can evolve extreme trait combinations with equal probability as central trait values. According to this null model, beak morphospace would approximate a cube. To simulate this, we sampled uniformly spaced values within the observed bounds of each trait axis independently. Null model 2 also assumes that each beak dimension varies independently, but that natural selection limits the exploration of extreme trait combinations, resulting in an approximately spherical morphospace. We implemented two variants of Null model 2. Null model 2.1 maintains the observed distribution of values along each trait dimension but randomly shuffles these independently across species, thus removing any coordination among trait dimensions. Null model 2.2 assumes a uniform distribution of values along each trait dimension but constrained these values to occur within a sphere of radius equal to half the mean range occupied by each trait dimension. The difference in expected volume and beak size variation between Null model 2.1 and Null model 2.2 was minor (**Supplementary Figure 1**), indicating that the distribution of values within the observed range of each trait dimension has a relatively minor effect on expected morphospace occupancy.

### Archetypal analysis of beak and body shape

To formally describe the shape of beak morphospace defined by PC2 and PC3 we applied two different approaches. First, we used an archetypal analysis implemented in the R package ‘archetypes*’* (69). This approach aims to identify irregular polygons described by k vertices or ‘archetypes’ such that the distribution of species trait values can be well represented as convex combinations of these archetypes. For k vertices, the algorithm randomly selects an initial position for each vertex and then iteratively explores different vertex positions, trying to minimize the residual sum of squares (RSS) between the position of each species in trait space and the boundaries of the polygon. The original archetype algorithm (69), requires these vertices to lie on the boundary of the convex hull enclosing the observations, making this method sensitive to extreme observations. Because we were interested in identifying the polygon shape that best describes the majority of bird forms, we used a ‘robust archetype’ analysis that downweights the importance of extreme values (70). We compared the fit of polygons with different numbers of vertices, from k=3 to 7. Because the procedure for optimising polygon fit is stochastic, for each value of k we repeated the analysis 100 times and from across the replicates calculated the median and 95% CI in RSS.

The best fitting polygons identified through an archetype analysis can take irregular forms so that shapes with more vertices will always describe the observed data better. For example, a polygon with 4 vertices can be constructed that will more closely fit the observed boundaries of morphospace, even if this shape is largely triangular in form (i.e. two of the vertices may be close together). We therefore developed a new shape fitting algorithm based on regular polygons, with edges of equal length. We systematically explored polygons from 3 to 7 vertices and finally a polygon with *n* = 360 vertices approximating a circle. A circle is expected either when there are either no constraints on morphospace or when there a very large number of trade-offs between trait axes. Our algorithm systematically varied polygon position, area and angle of rotation, allowing us to identify the polygon where the distance between the boundary of the polygon and the boundary of observed morphospace was minimised The boundary of observed morphospace was calculated using multivariate kernel density estimation in the R package ‘ks’ (71). The kernel was a multivariate normal distribution for each species centred on the trait values for each species with the optimal bandwidth selected using the *Hpi* function. We used the contour containing 90% of the total density of species as the boundary of empirical morphospace. The use of the 90% contour avoids estimates of morphospace shape being dominated by the small number of species with extreme traits. Because of the greater flexibility (i.e. allowing irregular shapes) we used the ‘robust archetype’ analysis to identify the location of archetypes and used our new regular polygon fitting approach, which is more conservative (i.e. is more likely to penalise shapes with many vertices), to confirm the number of vertices required to describe morphospace (70).

### The correspondence between archetypes and functional objectives

We visualised how feeding techniques and functional objectives are distributed throughout morphospace by overlaying a regular grid and then summing the relative % score of each technique/objective across the species within each cell. Because some techniques/objectives are relatively more commonly used than others (e.g. there are many more terrestrial than aquatic species), we quantified the proportional importance of each technique/objective in a cell weighted by the total % use of each technique/objective across all species. Thus, our maps showing techniques/objectives across morphospace indicate regions where each technique/objective is relatively highly represented accounting for its overall prevalence across birds (**Figure 2**).

To formally test whether specialization in a functional objective is higher closer to the vertices of the Pareto front we used beta regression. For each focal archetype, we rescaled the distance of species from that archetype to between 0.001 and 0.999. We then estimated the slope of the relationship between distance as the response variable and the % use of that functional objective as the predictor. We include a quadratic predictor term in the model to allow for non-linear responses (**Supplementary Figure 4, Supplementary Table 8**).

Our analysis is based on functional objectives scored using our understanding of the biomechanical requirements of different feeding techniques rather than actual measurements of the resource acquisition and processing efficiency of different beak shapes. To test that our interpretation of beak functional objectives is robust, we obtained independent data on bite force (*n* = 77 species, (36)) and bite speed (*n* = 18 species, (37)) derived from experiments across multiple species. We fit a linear model to estimate the slope of the relationship between bite force (log-transformed) or bite speed as response variables, and beak size (β_PC1_) and distance to Archetype 1 (β_dA1_) as predictors. Archetype 1 is the archetype that corresponds to high strength capacity in our dataset, and is thus expected to correspond to a high bite force but slow closing velocity. As expected, bite force increased with beak size (β_PC1 =_ 1.13 ± 0.98, p<0.001) and decreased with the distance from Archetype 1 (β_dA1_ = −0.40 ± 0.08, p<0.001) (**Supplementary Figure 5a**). Together, beak size and proximity to Archetype 1 explain a substantial proportion of the variation in bite force across species (R^2^ = 0.66). Bite speed is independent of beak size (p>0.05) and increases strongly with distance from Archetype 1 (β_dA1_ = 0.19 ± 0.04, p<0.001) (**Supplementary Figure 5b**). Distance from Archetype 1 explains 61% of the variation in bite speed. Thus, quantitative data on beak biomechanics support our interpretation for the trade-offs between strength capacity and resources acquisition (via reach or engulfment capacity) in constraining avian beak morphospace.

### Directionality in trait evolution

To explore the directionality of beak and body shape evolution, for each morphospace axis (i.e., PC2 and PC3 of beak and body) we performed an ancestral trait reconstruction using the *fastAnc* function from the ‘phytools’ R package (72). Because trait reconstructions can be sensitive to tree topology, we run the reconstruction analysis only on trees that include species with genetic data (n = 6670 species). We first used the inferred ancestral values to calculate the overall global direction of trait evolution across all phylogenetic branches, and specifically whether this was ‘inward’ towards the centroid of morphospace or ‘outward’, away from the centroid. To do this, we calculated the difference in the distance from the centroid between each ancestor and each of its two descendants, and report the ratio of inward and outward branches. We next calculated how the direction of trait evolution varies locally across morphospace by overlaying a regular grid and identifying the phylogenetic branches intersecting each grid cell based on the inferred morphological position of the ancestor and descendent node. We then calculated the circular mean and variance (V_obs_) in the angle of trait evolution.

We compared the inferred directionality of trait evolution at both global (i.e. across the entire tree) and local (i.e. within grid cells) scales to that expected under a null model of random trait evolution. We simulated traits under a Brownian motion model using the function *fitBM* from the R package ‘phytools’, setting the rate of trait evolution sigma *σ* = 1 and ancestral trait value of 0. As with our observed trait data, we performed an ancestral reconstruction on these simulated values. To test for non-random directionality globally, we estimated the null ratio of outward and inward branch trajectories. We repeated our calculation of the observed and simulated directionality ratio across the same 100 phylogenetic trees and report the 95% CI in the proportion of lineages showing outward directionality. The same analysis was performed using the beak shape axes and the body shape axes independently.

To test for non-random directionality locally, we used an alternative procedure which is necessary because in any given null model simulation a grid cell may fail to contain any inferred branches or contain far more or fewer than is inferred for the observed data due to the model of random trait evolution. Our null model of local directionality therefore used the following steps. First, for each simulation run we calculated the variance in the inferred direction of evolution (V_sim_) for each grid cell. Second, we plotted how V_sim_ varies according to the number of branches intersecting a cell (NB_sim_). For this, we pooled values of V_sim_ and NB_sim_ across simulation runs. Third, we identified the boundary describing the minimum V_sim_ expected for a given value of NB_sim_ using a quantile regression and the 5% quantile (i.e. 1-tailed test). Fourth, given the number of branches in our empirical data inferred to intersect each cell (NB_obs_) we identified values of V_obs_ that were lower than expected under the null model. Here, a significantly lower variance indicates that co-occurring branches tend to evolve in a more consistent direction than expected under a null model of Brownian trait evolution.

### Mapping extinction risk and speciation rates across morphospace

To explore how different regions of morphospace are associated with variation in extinction risk, we used species threat status from the IUCN Red List (version 2022-1)(73). We used a binary classification, considering species classified as LC as not threatened (0) and the rest (NT, VU, EN, CR, EX) as threatened (1). Species classified as Data deficient (DD) were excluded (*n* = 38). To explore how different regions of morphospace are associated with variation in speciation rate, we used the DR metric (65). The DR metric is the inverse of the equal splits (ES) metric of evolutionary isolation (74) and although originally considered a measure of net diversification, it better approximates speciation rate (75). We used the function “evol.distinct” from the R package ‘picante’ (76) to calculate the DR of each species across the complete BirdTree phylogeny. We then identified those species in the top DR quartile as having high speciation rates (1), with the remaining species scored as having relatively low (0) speciation rates. Then, extinction risk (1= Threatened / 0 = Non-threatened) and speciation rate (high=1 vs low =0) were modelled as a function of the position of species in both beak and body shape morphospace, using a generalized additive model (GAM) with a binomial error distribution implemented in the R package ‘mgcv’ (77). The position of species in either beak or body shape morphospace (i.e. PC2 and PC3) were included as predictors and used to predict the relative incidence of lineages with high extinction risk and high speciation rates across morphospace.

### Testing the effect of morphological position on extinction and speciation

To test the effect of biophysical specialization on the risk of extinction and on speciation rates, we fit a species level Bayesian linear mixed models accounting for the phylogenetic non-independence of species, using the R package ‘MCMCglmm’ (78). We modelled extinction risk (1= Threatened / 0 = Non-threatened) as a binary-response with a probit link (family ordinal). For DR (log-transformed), we used a normal distribution (family gaussian). In both models, the predictors were the degree of biophysical specialization in beak and body shape, calculated as the distance to the nearest beak and body shape archetype, respectively. We also included other factors expected to influence extinction risk and speciation rate, including degree of insularity, habitat breadth and generation length (log-transformed). Body size was not included as it was highly correlated with generation length (Pearson’s r=0.84) and might cause co-linearity problems. The degree of insularity was assessed from the BirdLife International distribution maps (79). We classified each species as insular (value of 1) when occurring year round on oceanic islands that do not reconnect to the continent when sea levels changed during glacial periods (considering the minimum level of 120 m below current level) (80). Species that occur on continental or land bridge islands, that do reconnect with continents were given a value of 0.5, whereas species that occur on continents were given a value of 0. Habitat breadth was obtained from Ducatez et al. (81), which employs a multiplicative beta diversity index derived from the presence/absence of species along 82 different habitat subtypes from the IUCN (73). Generation length was obtained from Bird et al. (82). Each model was run for 110,000 iterations (with a 10,000 burn-in and 100 of thinning interval). The thinning interval was set to 100, resulting in a posterior distribution of 1,000 samples, and sufficient so ensure that the autocorrelation of samples was <0.1. The models include the phylogenetic effects as a random factor, using a maximum clade credibility tree (MCC) from the posterior sample of full trees.

