## Supplementary tables and figures for "Biophysical constraints on avian adaptation and diversification"

### This file contains:

**Supplementary Tables (1-8)**

**Supplementary Figures (1-11)**

**Supplementary Table 1. Loadings of beak dimensions of each PC axis.** The % of total variance accounted for by each PC axis is indicated.

| Trait | PC1 (85.8 %) | PC2 (12.0%) | PC3 (2.2%) |
|---|---|---|---|
| Beak length | 0.538 | 0.843 | -0.023 |
| Beak width | 0.595 | -0.399 | -0.698 |
| Beak depth | 0.597 | -0.362 | 0.716 |

**Supplementary Table 2.** Scoring of beak functional objectives across feeding techniques.

| Feeding technique | Strength | Engulfment | Reach |
|---|---|---|---|
| Tearing | 1 | 0 | 0 |
| Crushing | 1 | 0 | 0 |
| Engulfing | 0 | 1 | 0 |
| Filtering | 0 | 0.75 | 0.25 |
| Sweeping | 0 | 0.25 | 0.75 |
| Grazing | 0 | 0.75 | 0.25 |
| Hammering | 0.5 | 0 | 0.5 |
| Plucking | 0.33 | 0.33 | 0.34 |
| Probing | 0 | 0 | 1 |
| Spearing | 0.25 | 0 | 0.75 |

**Supplementary Table 3. Loadings for body traits on the first three PC analysis**. The % of total variance accounted for by each PC axis is indicated.

| Trait | PC1 (74.9 %) | PC2 (16.0%) | PC3 (8.3%) |
|---|---|---|---|
| Tarsus length | 0.458 | 0.659 | -0.516 |
| Tail length | 0.498 | 0.207 | 0.820 |
| Wing length | 0.568 | -0.128 | -0.124 |
| Kipp’s distance | 0.467 | -0.711 | -0.217 |

**Supplementary Table 4.** Scoring of body functional objectives across foraging niches.

| Foraging technique | Aerial | Terrestrial | Aquatic |
|---|---|---|---|
| Invertivore aerial screening | 1 | 0 | 0 |
| Invertivore sally-to-air | 1 | 0 | 0 |
| Invertivore sally-to-surface | 0.5 | 0.5 | 0 |
| Invertivore sally-to-ground | 0.5 | 0.5 | 0 |
| Invertivore vertical substrate | 0.5 | 0.5 | 0 |
| Invertivore glean elevated | 0.5 | 0.5 | 0 |
| Invertivore glean ground | 0 | 1 | 0 |
| Frugivore aerial | 0.5 | 0.5 | 0 |
| Frugivore glean | 0.5 | 0.5 | 0 |
| Frugivore ground | 0 | 1 | 0 |
| Nectarivore aerial | 1 | 0 | 0 |
| Nectarivore glean | 0.5 | 0.5 | 0 |
| Granivore elevated | 0.5 | 0.5 | 0 |
| Granivore ground | 0 | 1 | 0 |
| Herbivore elevated | 0.5 | 0.5 | 0 |
| Herbivore ground | 0 | 1 | 0 |
| Vertivore aerial screening | 1 | 0 | 0 |
| Vertivore air-to-surface | 0.5 | 0.5 | 0 |
| Vertivore perch | 0.5 | 0.5 | 0 |
| Vertivore glean elevated | 0.5 | 0.5 | 0 |
| Vertivore glean ground | 0 | 1 | 0 |
| Carrion aquatic | 0 | 0 | 1 |
| Carrion ground | 0 | 1 | 0 |
| Herbivore aquatic ground | 0 | 0.5 | 0.5 |
| Herbivore aquatic surface | 0 | 0 | 1 |
| Herbivore aquatic dive | 0 | 0 | 1 |
| Aquatic predator ground | 0 | 0.25 | 0.75 |
| Aquatic predator perch | 0.5 | 0 | 0.5 |
| Aquatic predator air | 0.5 | 0 | 0.5 |
| Aquatic predator plunge | 0.5 | 0 | 0.5 |
| Aquatic predator surface | 0 | 0 | 1 |
| Aquatic predator dive | 0 | 0 | 1 |

**Supplementary Table 5.** Bayesian phylogenetic mixed effect models, with extinction risk (model 1) and diversification rate (model 2) as a function of distance to closest body and beak archetype, as well as other confounding factors, including generation length, habitat breadth and insularity (*n* = 9, 873 species).

| | **Post. mean** | **l-95% C.I.** | **u-95% C.I.** | **pMCMC** |
|---|---|---|---|---|
| **Model 1 (Extinction risk)** | | | | |
| (Intercept) | -2.018 | -3.100 | -0.960 | 0.002 |
| Dist. closest beak archetype | -0.021 | -0.128 | 0.087 | 0.682 |
| Dist. closest body archetype | -0.134 | -0.247 | -0.026 | 0.024 |
| Log (Generation length) | 0.862 | 0.673 | 1.082 | <0.001 |
| Habitat breadth | -0.152 | -0.194 | -0.113 | <0.001 |
| Insularity index | 1.521 | 1.370 | 1.667 | <0.001 |
| **Model 2 (Diversification rate)** | | | | |
| (Intercept) | -2.813 | -3.236 | -2.380 | <0.001 |
| Dist. closest beak archetype | -0.004 | -0.005 | 0.040 | 0.902 |
| Dist. closest body archetype | 0.016 | -0.032 | 0.065 | 0.556 |
| Log (Generation length) | -0.009 | -0.081 | 0.086 | 0.830 |
| Habitat breadth | 0.034 | 0.017 | 0.051 | <0.001 |
| Insularity index | 0.174 | 0.109 | 0.242 | <0.001 |

**Supplementary Table 6. Foraging niche (*n* = 32)**. The number of species specializing on each niche (>60% of use) is indicated. Species that do not use any of the categories over 60% are considered generalists (*n* = 3441 species).

| Foraging niche | Description | number of specialist species |
|---|---|---|
| Invertivore aerial screening | Species capturing flying invertebrates on the wing (e.g. swallows, swifts). Often described as 'screening' or 'hawking'. In contrast to 'Invertivore Sally air', characterized by continuous and extended flight with multiple items captured before landing. | 270 |
| Invertivore sally-to-air | Species capturing flying invertebrates in mid-air, with the attack starting from a perch (i.e. branch, rock, fence post, telegraph wire, etc.) and then returning to a perch (e.g. jacamars, kingbirds, etc). 'Hawking' will sometimes refer to this category, but the key distinguishing feature is that only a single prey item is captured before returning to a perch. | 278 |
| Invertivore sally-to-surface | Species capturing invertebrates (including arachnids, worms, molluscs, etc.) attached to the substrate (e.g. leaves, twigs, branches, rock faces, etc) following an aerial attack manoeuvre (e.g. flight, pounce, jump, hover). | 278 |
| Invertivore sally-to-ground | Species capturing invertebrates on the ground following an aerial attack manoeuvre (e.g. flying, gliding, dropping or pouncing) (e.g. chats, shrikes, kiskadee etc). The aerial manoeuvre may be followed by brief hopping toward prey (e.g. terns). | 152 |
| Invertivore vertical substrate | Species capturing invertebrates attached to or concealed within large branches and trunks of trees (e.g. woodpeckers, treecreepers, woodcreepers, wallcreepers, nuthatches, sittelas, nuthatches, vangas, etc., including honeyguides). This is distinguished from 'Invertivore Glean arboreal' by at least one criterion. First, the species employs specialized methods for moving over surfaces which are often, but not always vertical, vertical and too large to be gripped by the closed foot (including creeping, climbing or scaling). Second, the species extracts prey from in/under the bark using specialized methods (including hammering, probing or chiselling). Also includes species capturing insects from rock and cliff-faces (though not just on boulders), habitually perching on or clinging to large mammals and species that feed on honey and beeswax. | 282 |
| Invertivore glean elevated | Species capturing invertebrates attached to the substrate (e.g. leaves, twigs, branches, grass, bamboo, stems, hanging dead-leaves [not dead leaves on the ground] etc). No aerial attack manoeuvre is involved. | 1547 |
| Invertivore glean ground | Species capturing invertebrates on the ground. In contrast to 'Invertivore Sally ground', the search and attack manoeuvres take place on the ground (e.g. thrushes). This includes species standing on the ground and gleaning insects from vegetation (e.g. tinamous or larks) but excludes species that jump or sally upwards to capture prey from vegetation ('Invertivore Sally surface') or the air ('Invertivore Sally air'). The ground is dry and thus excludes aquatic habitats (e.g. beaches, estuaries, wetlands, marshes ['Aquatic predator Ground' ]). | 811 |
| Vertivore aerial screening | Species captures vertebrate prey during flight. Both predator and prey are in flight (e.g. peregrine, hobby, falcon) | 12 |
| Vertivore air-to-surface | Vertivore Air to surface (n = 54) – species captures prey on branches or the ground by diving from the air, usually after circling or hovering in flight. Includes quartering flight (e.g. kestrels, kites, some owls). | 44 |
| Vertivore perch | Species captures prey on branches or the ground by diving from a perch (e.g. many owls, eagles). | 155 |
| Vertivore glean elevated | Species capturing prey from foliage, branches, epiphytes, cavities, bark or other arboreal substrate while perched on the substrate. There is no flight attack involved. This includes eating bird chicks from arboreal nests and drinking blood while perched on mammals (e.g. oxpeckers). | 4 |
| Vertivore Glean Ground | Species capturing prey on the ground, including eggs in ground nests, while they themselves are also walking or running on the ground (e.g. secretary bird, seriemas, ground hornbills). | 3 |
| Carrion aquatic | Species eating carrion (dead animal or fish remains) | 0 |
| Carrion ground | Species eating carrion (dead animal or fish remains) on the ground (e.g. vultures). | 22 |
| Frugivore aerial | Species foraging on fruits in flight, including those that hover to pluck fruit from bushes and trees (e.g. oilbird, some manakins). | 79 |
| Frugivore glean | Species foraging on fruits while perched (not in flight) above ground and plucking fruits from vegetation (e.g. toucans, hornbills). | 855 |
| Frugivore ground | Species foraging on fruits lying on the ground (e.g. trumpeters). | 18 |

| | | |
|---|---|---|
| Nectarivore aerial | Species feeding on nectar or other plant exudates (e.g. sap) while in flight (e.g. hummingbirds). | 308 |
| Nectarivore glean | Species feeding on nectar or other plant exudates (e.g. sap) while perched, including nectar predators that pierce corollas (e.g. sunbirds, flowerpiercers). Species feeding on honey (e.g. honeyguides) included under 'Invertivore Bark'. | 148 |
| Granivore arboreal | Species foraging on seeds, grains and nuts taken from vegetation (e.g. trees, grass stems) while perched (e.g. seedeaters, finches). | 137 |
| Granivore ground | Species foraging on fallen seeds, grains and nuts collected from the ground (e.g. partridges, pheasants, finches). | 320 |
| Herbivore elevated | Species foraging on leaves, buds, blossom, or other vegetation (except fruit, seeds and nectar). The food is taken from above ground, generally while the species is perching on branches or other stems. Generally, a small part of diet, except for the hoatzin, plantcutters. | 12 |
| Herbivore ground | Species foraging on grass, leaves, buds, blossom, or other vegetation (not fruit or seeds) taken while the species is on the ground (e.g. geese). The vegetation may itself be off the ground. | 63 |
| Aquatic predator ground | Species capturing invertebrates or vertebrates while standing in aquatic habitats (including beaches, estuaries, wetlands and marshes) (e.g. storks, herons, shorebirds). Prey may be captured on the ground or on/under water. This category includes species capturing aquatic prey (e.g. fish) or terrestrial prey in aquatic habitats (e.g. grasshopper). | 298 |
| Aquatic predator perch | Species capturing invertebrates or vertebrates on/under water following a direct attack flight from a perch (e.g. kingfisher). | 40 |
| Aquatic predator air | Species capturing invertebrates or vertebrates on/under water during continuous flight (including dipping, hovering, pattering, snatching). In contrast to 'Aquatic predator Perch', prey item is identified while flying (not from perch). The predators body may partially submerge but does not plunge beneath the surface (see 'Aquatic predator Plunge'). Includes kleptoparasitic species capturing fish by chasing other piscivores and forcing them to regurgitate (e.g. skuas, frigatebirds). | 79 |
| Aquatic predator plunge | Species capturing invertebrates or vertebrates by plunging under water following continuous flight. The predators body submerges entirely beneath the surface, with the prey captured either by the momentum of the plunge or following propelled swimming. | 46 |
| Aquatic predator surface | Species capturing invertebrates or vertebrates on/under water whilst swimming on the water surface. In contrast to 'Aquatic predator Perch' or 'Aquatic predator Aerial' there is no direct attack flight. The species may dip under the water but, in contrast to 'Aquatic predator Dive', contact with the surface is maintained. | 54 |
| Aquatic predator dive | Species capturing invertebrates or vertebrates under water by diving from the surface (not the air, see 'Aquatic predator Perch' and 'Aquatic predator Plunge'). | 127 |
| Herbivore aquatic ground | Species foraging on aquatic vegetation (including seeds) either below or above the water surface (algae, pondweed, waterside vegetation). The species collects vegetation while under water, sitting on the water surface or wading. | 8 |
| Herbivore aquatic surface | Species foraging on aquatic vegetation (including seeds) on/under water whilst swimming on the water surface. The species may dip under the water but, in contrast to 'Herbivore (A) dive', contact with the surface is maintained. | 17 |
| Herbivore aquatic dive | Species foraging on aquatic vegetation (including seeds) under water by diving from the surface. | 4 |

**Supplementary Table 7. Beak feeding techniques (*n* = 10).** The number of species specializing on each technique (>60% of use) is indicated. Species that do not use any of the categories over 60% are considered generalists (*n* = 1290 species).

| Feeding technique | Description | number of specialist species |
|---|---|---|
| Tearing | Beaks are used to pull apart pieces of living or dead prey. Examples include raptors, vultures, skuas or gulls pulling off flesh from vertebrate or large invertebrate (e.g. squid) prey. Also includes species tearing leaves from trees (e.g. hoatzin), cutting reeds (e.g. Parrotbills) and using the beak to dig (e.g. *Cacatua tenuirostris*). | 393 |
| Crushing | Beaks are used to break open hard resources such as shells, seeds, or nuts by squeezing and force of pressure (e.g. parrots and various kinds of finches). Crush also includes other high bite-force actions such as mashing and cutting hard substrates (for example *Catamblyrhynchus diadema* cuts through tough bamboo stems with its beak). | 167 |
| Engulfing | The beak is designed to maximize surface area or volume that is captured in a snatching, sweeping or lunging manoeuvre. Examples include nightjars, swifts and swallows catching prey in the air and pelicans catching prey under water. Flycatchers or nightjars that capture prey in continuous or sallying flights, either to the air or to substrates, are classified as engulfers if the attack manoeuvre is imprecise. That is, the prey is not trapped precisely between the mandibles but instead engulfed within the mouth, often because the predator is hunting prey that is obscured by darkness, deep shade, or foliage. | 852 |
| Filtering | Use of the beak and its internal structures as a fine grill to filter water and obtain plankton or small invertebrates. Examples include flamingos, swans and some ducks. | 121 |
| Sweeping | Use of the beak as a touch sensitive tool to detect prey at or beneath the surface of water. Examples include spoonbills, avocets, and skimmers. | 26 |
| Grazing | The beak is used to feed on soft plant material such as grass or aquatic plants (e.g. some ducks and rails). Grazing of vegetation on level ground or under water is like pecking of seeds in as much as it takes little bite force and generally involves little or no reaching away from the body. | 26 |
| Hammering | Beaks are used to reach resources embedded within wood using hard impacts for chipping or chiselling the substrate. Examples include woodpeckers chipping dead wood to reach larvae, and species using the beak as a hammer to break open seeds, shells or carapaces (e.g. oystercatchers breaking open mussels or tits opening sunflower seeds), or using bill to prise open fruit (e.g. Sharpbill) | 29 |
| Plucking | Involves a precise manoeuvre to obtain a food item from air, water or surface. Examples include: insectivores (e.g. warblers) reaching to snatch insects from foliage; flycatchers sallying to snatch insects from the underside of leaves (see also engulfing); shorebirds (e.g. plovers and most sandpipers) picking up invertebrates or vertebrates from the surface of (or very near the surface of) sediments, frugivores reaching to pick fruit & berries from vegetation and vertivores such as forest kingfishers, storks, gulls and albatrosses flying or reaching down to snatch items from the ground or on/near the water's surface. Within the plucking category, we also include pecking, where the beak is used to pick food items from a flat substrate. Examples include many finches, pigeons or gamebirds softly pecking for seeds on the ground | 6402 |
| Probing | Involves the relatively slow or gentle insertion of the beak into a crevice or beneath the ground surface to extract food. Examples include invertivores inserting beaks into narrow apertures (e.g. treecreepers, wood-hoopoes and scythebills looking for prey under tree bark or in openings in wood or bamboo), or inserting beaks into soft sediment (e.g. curlews, godwits, and some sandpipers in wetland habitats), as well as nectarivores probing to reach nectar in flowers (e.g. hummingbirds and sunbirds). | 602 |
| Spearing | Involves the use of the beak as a dagger to stab prey. Examples include anhingas and herons catching fish or rodents and oystercatchers stabbing bivalves. | 4 |

**Supplementary Table 8.** Relationship between distance from each archetype and specialisation in the corresponding functional objective ($n$ = 9, 908 species).

| Archetype | Intercept | Slope | Quadratic |
|---|---|---|---|
| Engulfment (A3) | 0.22 ± 0.01*** | -0.72 ± 0.05*** | -0.38 ± 0.05*** |
| Strength (A1) | 0.29 ± 0.01*** | -2.79 ± 0.05*** | 1.25 ± 0.06*** |
| Reach (A2) | 0.15 ± 0.01*** | -1.57 ± 0.05*** | 0.08 ± 0.06 |
| Engulfment (A3) | 0.22 ± 0.01*** | -0.72 ± 0.05*** | -0.38 ± 0.05*** |
| Aquatic (A1) | -0.58 ± 0*** | -2.08 ± 0.08*** | 1.51 ± 0.1*** |
| Terrestrial (A2) | 0.88 ± 0.01*** | -2.62 ± 0.05*** | 1.89 ± 0.05*** |
| Aerial (A3) | -0.62 ± 0.01*** | 0.15 ± 0.03*** | -1.16 ± 0.03*** |

***P<0.001, **P<0.01, *P<0.05

**Supplementary Figures (1-8)**

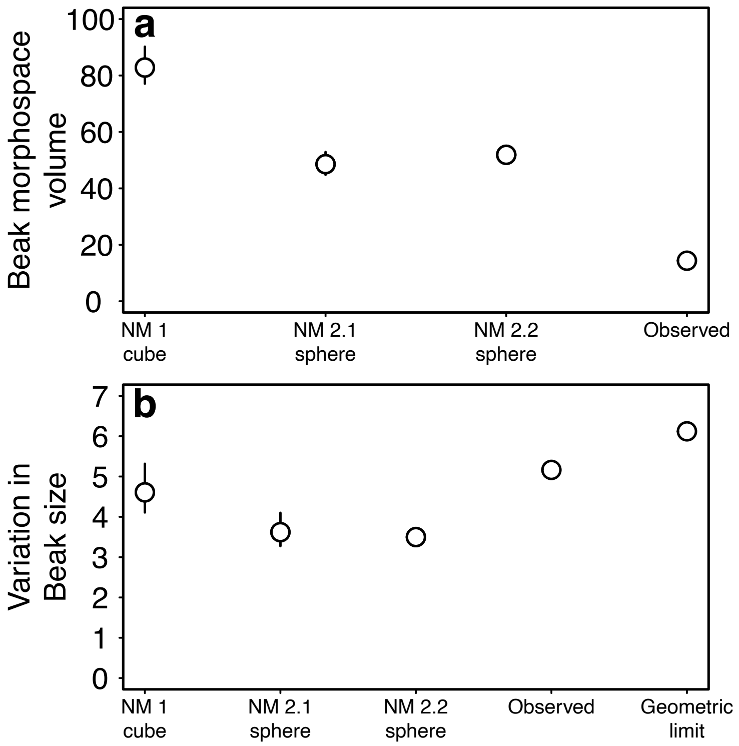


**Supplementary Figure 1. Null models of avian (a) beak morphospace volume and (b) variation in beak size across species (*n* = 9, 993 species)**. In (**a**) volume of beak morphospace is the minimum convex hull containing log-transformed beak length, width and depth measurements for observed data and null model (NM) simulations. In (**b**) beak size is calculated as the beak volume assuming beaks have a conical shape. Variation in beak size is the order of magnitude change from the smallest to largest beak. The geometric limit shows the maximum possible variation in beak size given the observed range of each trait dimension. Points and lines show the median and 95% C.I. respectively.

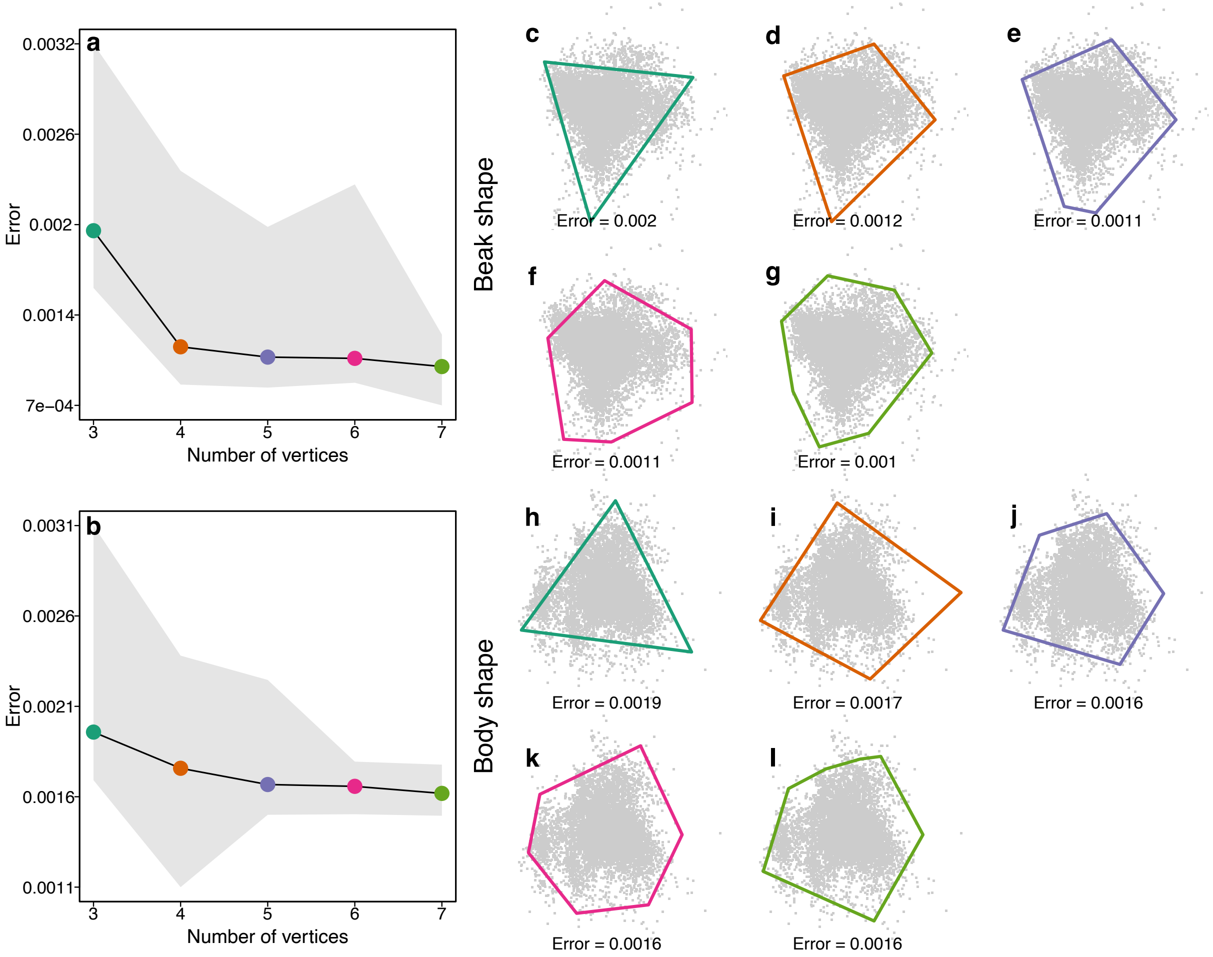


**Supplementary Figure 2. Best fitting irregular polygons describing beak (a) and body (b) shape morphospace (*n* = 9, 993 species).** (**a-b**) show the error (residual sum of squares) of polygons with different numbers of vertices. Lines and grey shading show the median and 95%CI across 100 replicate iterations. For each vertex number (*n* = 3-7), the best fitting polygons for beak (**c-g**) and body (**h-l**) shape morphospace are shown. The archetype algorithm rarely converged for polygons with more than 7 vertices and so results for more complex shapes are not shown.

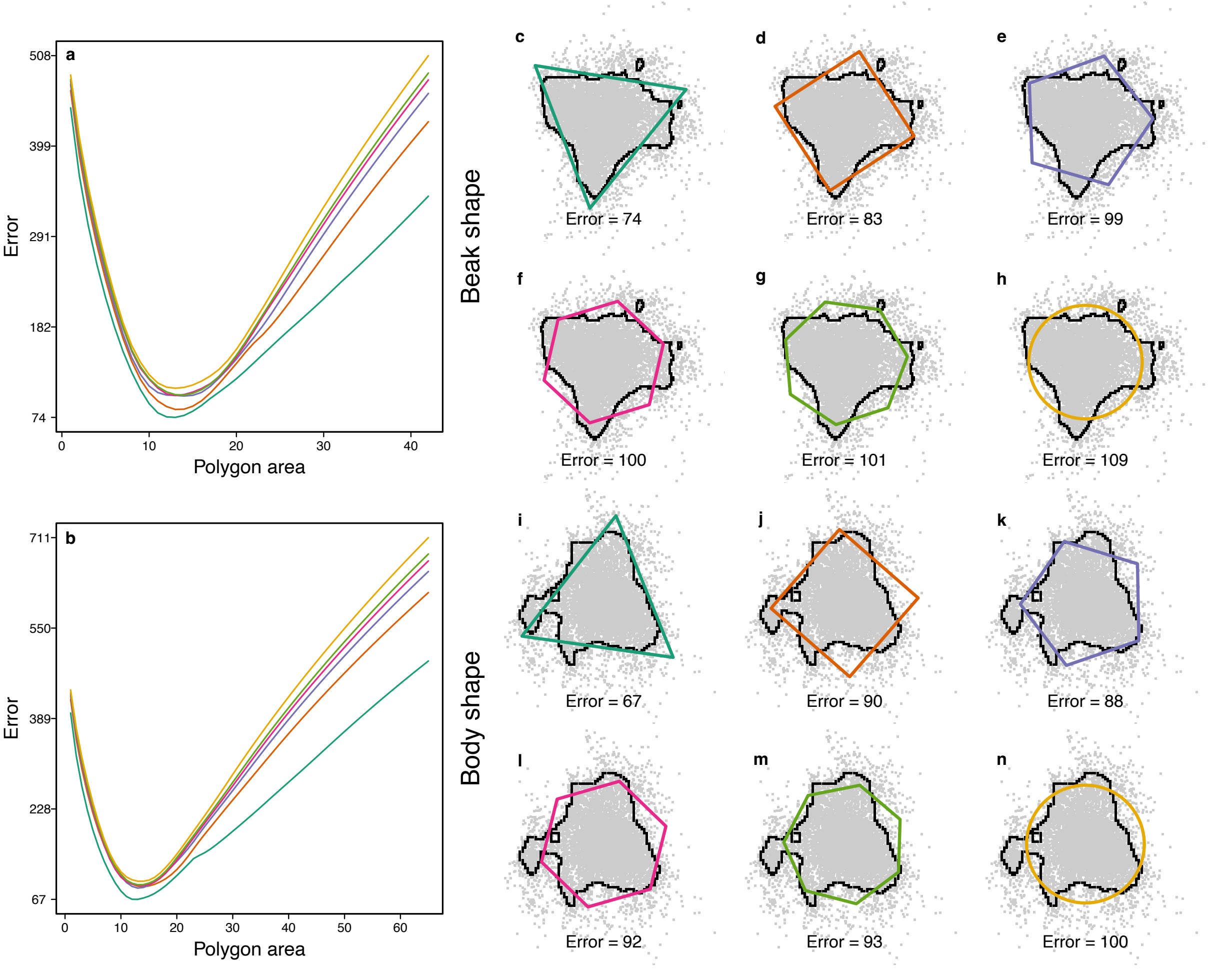


**Supplementary Figure 3. Best fitting regular polygons describing beak (a) and body (b) shape morphospace (*n* = 9, 993 species).** (**a-b**) show the error (sum of absolute distances between polygon and morphospace boundary) of polygons with different numbers of vertices and areas. The morphospace boundary is the irregular black border corresponding to the 90% density quantile of species trait values (grey points). For each vertex number (*n* = 3-7 and 360 vertices), the best fitting polygons for beak (**c-h**) and body (**i-n**) shape morphospace are shown.

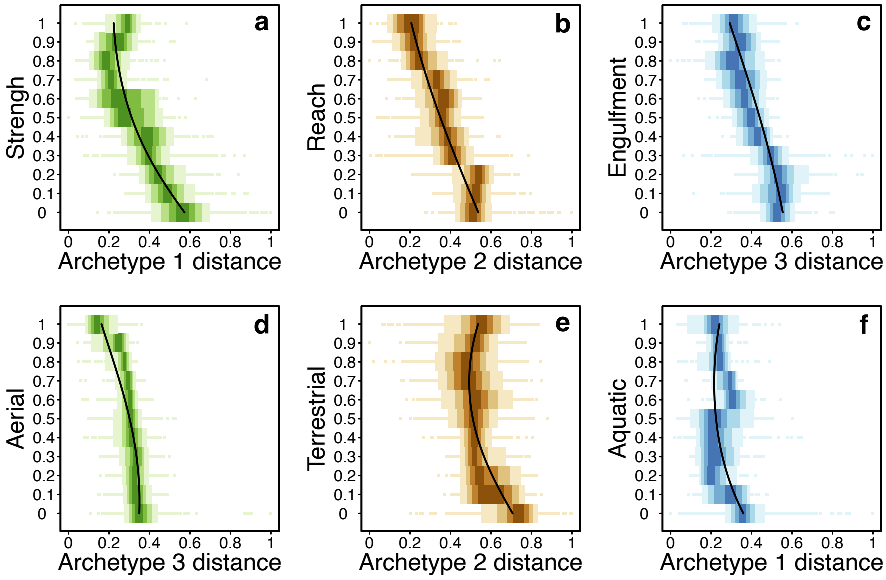


**Supplementary Figure 4. Specialisation on each functional objective task increases towards the respective archetype (i.e. vertex) of the triangular pareto.** Results are shown for beak (**a-c**) and body (**d-f**) shape morphospace ($n$ = 9, 993 species). Specialisation (y-axis) is the % importance of the (**a-c**) beak functional objective or (**d-f**) body functional objective for feeding. For each level of specialisation, box plots show the distribution of distances from the focal archetype (color bands of boxes show 10% quantiles). Fitted lines are from a quadratic beta-regression predicting distance as a function of specialisation on the respective functional objective. Distances to archetypes were rescaled between 0.01 and 0.99 for model fitting and plotting.

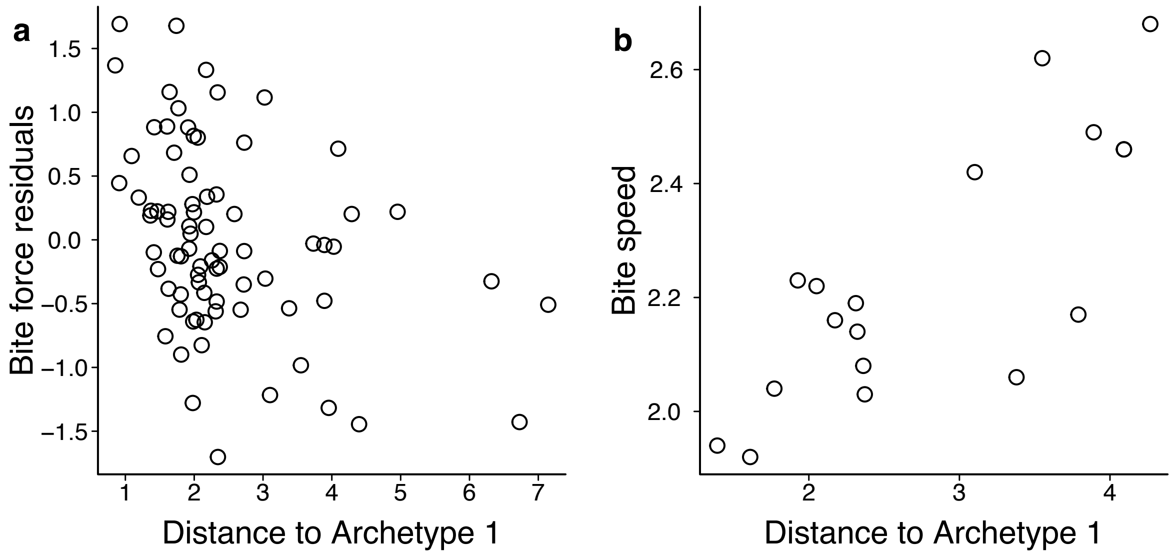


**Supplementary Figure 5. Beak shape and biomechanics.** (**a**) Bite force declines with distance from Archetype 1 ($n$ = 77 species), while (**b**) bite speed (i.e. beak closing velocity) increases with distance from Archetype 1 ($n$ = 18 species). In (**a**) bite force residuals are from a linear model predicting bite force (log-transformed) as a function of beak size (PC1).

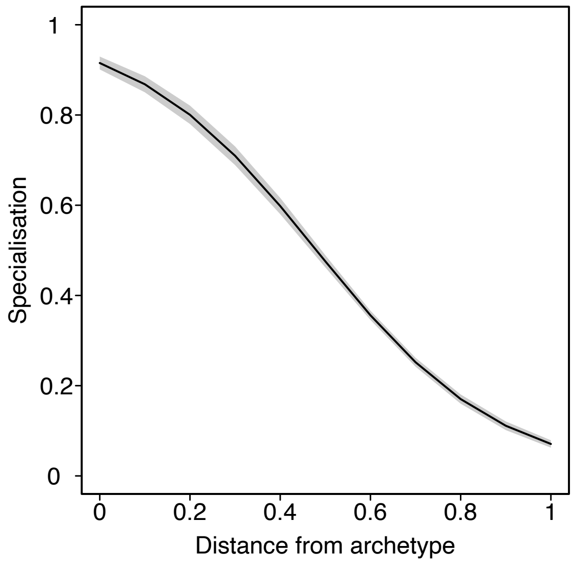


**Supplementary Figure 6. Specialisation in beak functional objectives increases towards the archetypes (i.e. vertices) of the triangular pareto front (*n* = 9, 993 species).** Distance is calculated to the nearest archetype and scaled between 0 and 1. Species were assigned as specialists if the importance of any functional objective to resource acquisition is ≥60%. The fitted relationship is from a generalised linear model with a binomial error distribution.

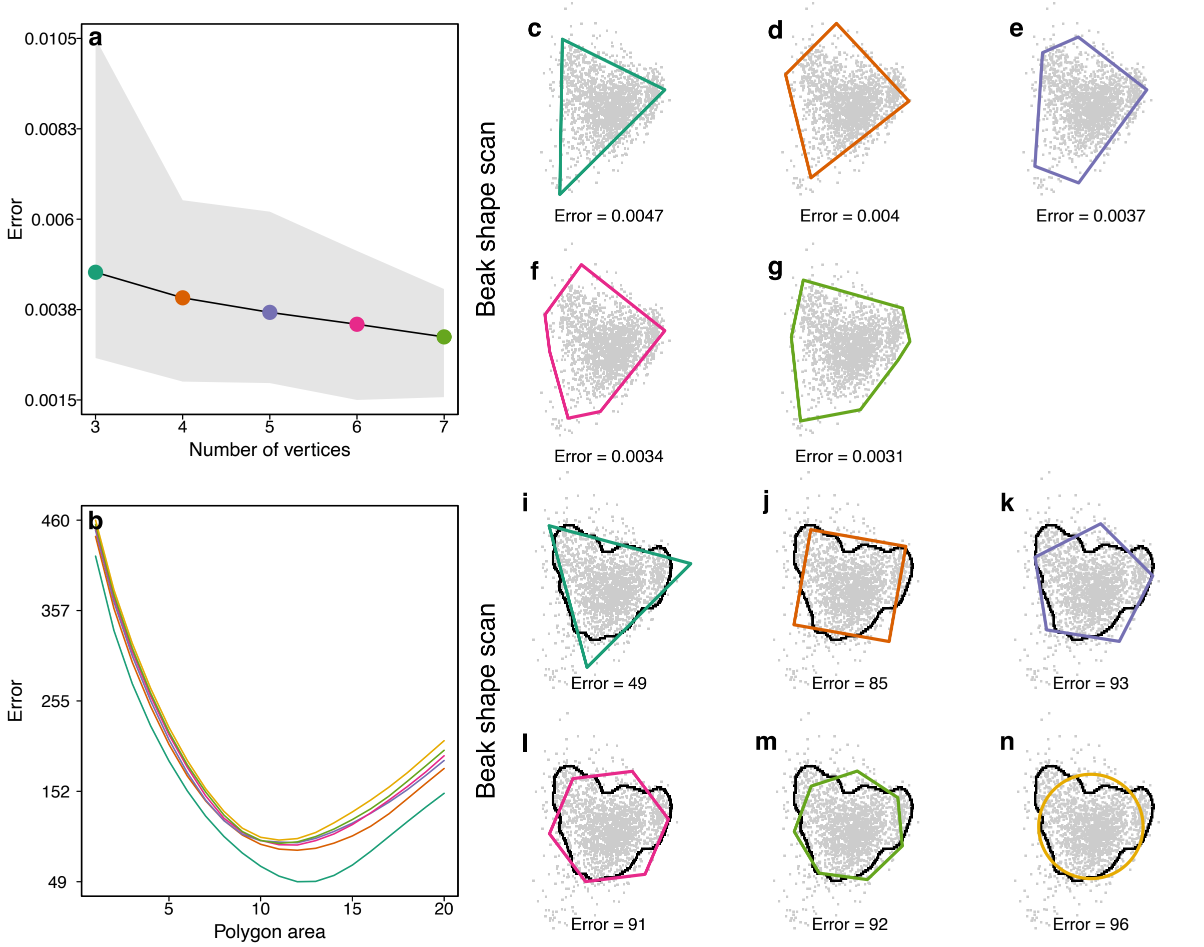


**Supplementary Figure 7. Best fitting irregular and regular polygons describing beak shape morphospace derived from 3D scans of the beak (*n* = 2, 028 species).** (**a-b**) show the error of polygons with different numbers of vertices. In (**a**) lines and grey shading show the median and 95%CI in the residual sum of squares across 100 replicate iterations. In (**b**) lines show the sum of absolute distances between the polygon and morphospace boundary for polygons with different numbers of vertices and areas. For each vertex number (*n* = 3-7 and 360 vertices), best fitting (**c-g**) irregular and (**i-n**) regular polygons are shown. In (**i-n**) the morphospace boundary is the irregular black border corresponding to the 90% density quantile of species trait values (grey points).

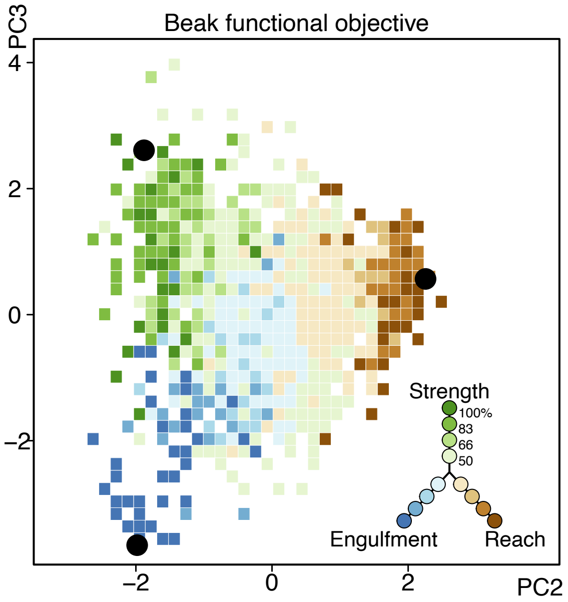


**Supplementary Figure 8. Beak shape morphospace derived from 3D scans (*n* = 2028 species).** Black points show the position of the 'archetypes' (i.e. vertices) defining the best fitting triangle for describing beak shape variation. Colors indicate the relative dominance of each of the three functional objectives (strength, reach and engulfment capacity).

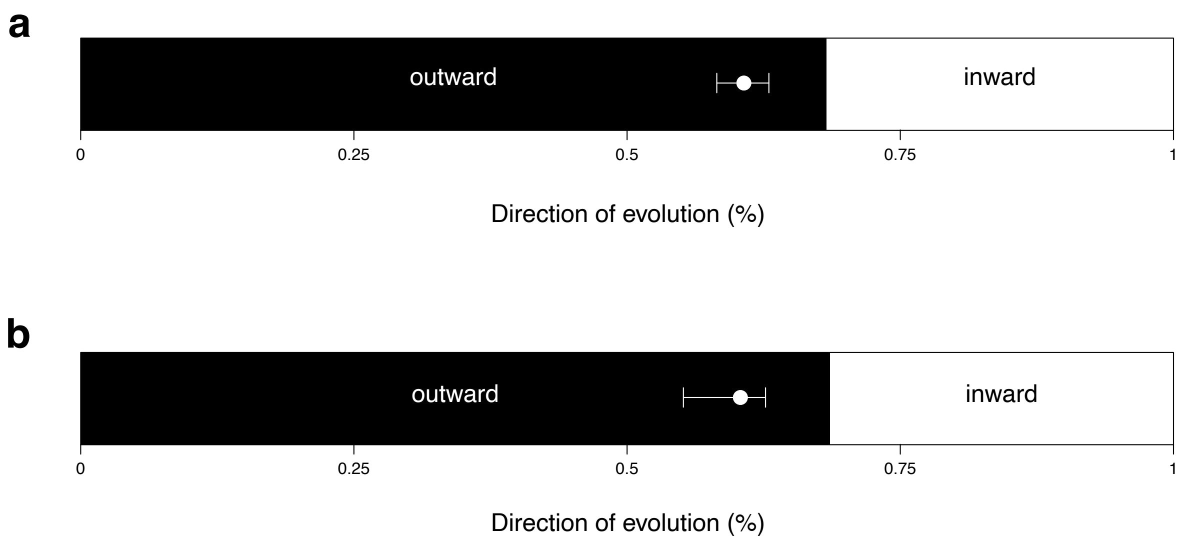


**Supplementary Figure 9. Directionality of beak and body shape evolution.** The proportion of lineages moving outwards towards the edge (black) or inwards towards the centre (white) of beak (**a**) and body (b) shape morphospace. The white bracket shows the null expectation (95% CI) under a model of random trait evolution.

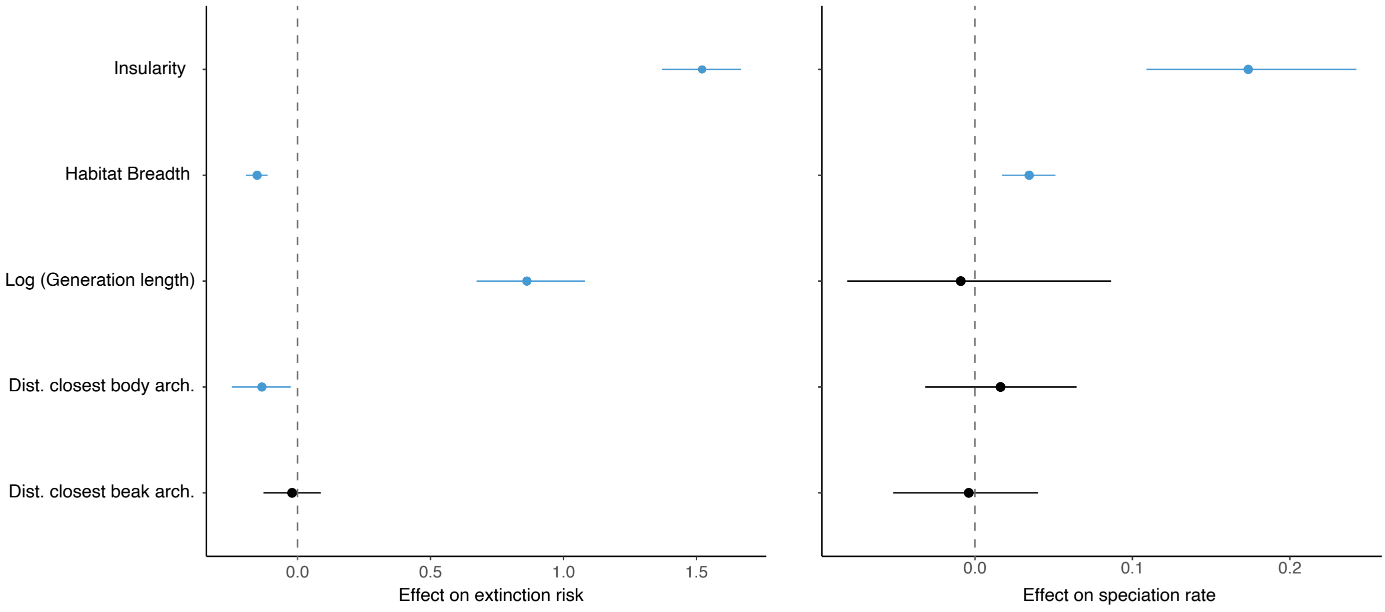


**Supplementary Figure 10. Effect of foraging specialization and other factors in speciation and extinction.** We show the effect of distance to closest beak and body archetypes and other factors (insularity, habitat breadth and generation length) on extinction risk (left panel) and speciation rates (right panel). For each factor, we show the distribution of effect sizes (mean and 95 % C.I.) from the posterior distribution of Bayesian phylogenetic mix models. Significant factors (not overlapping with zero) are colored in blue.

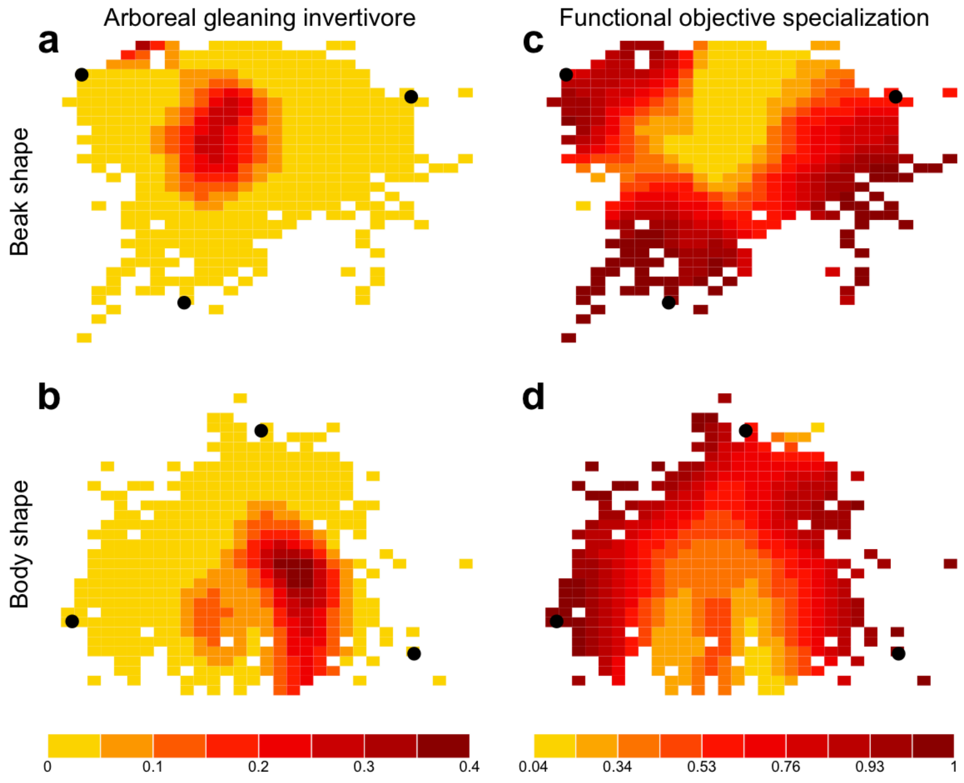


**Supplementary Figure 11. Specialization in arboreal invertebrate gleaning and functional objectives across beak and body shape morphospace.** Predicted probability that a species is a specialist arboreal invertebrate gleaner (listed as "Invertivore glean elevated" in **Supplementary Table 6**) across beak (**a**) and body (**b**) morphospace. Predicted probability that a species is specialized on a single (**c**) beak and (**d**) body functional objective across beak and body morphospace respectively.
